# Decoding dynamic visual scenes across the brain hierarchy

**DOI:** 10.1101/2024.06.24.600332

**Authors:** Ye Chen, Peter Beech, Ziwei Yin, Shanshan Jia, Jiayi Zhang, Zhaofei Yu, Jian K. Liu

**Affiliations:** School of Computer Science, Peking University, Beijing, China; Institute for Artificial Intelligence, Peking University, Beijing, China; School of Computing, University of Leeds, Leeds, UK; School of Computer Science, Centre for Human Brain Health, University of Birmingham, Birmingham, UK; Institutes of Brain Science, State Key Laboratory of Medical Neurobiology, MOE Frontiers Center for Brain Science and Institute for Medical and Engineering Innovation, Eye & ENT Hospital, Fudan University, Shanghai, China

## Abstract

Understanding the computational mechanisms that underlie the encoding and decoding of environmental stimuli is a paramount investigation within the domain of neuroscience. Central to this pursuit is the exploration of how the brain represents visual information across its hierarchical architecture. A prominent challenge resides in discerning the neural underpinnings of the processing of dynamic natural visual scenes. Although considerable research efforts have been made to characterize individual components of the visual pathway, a systematic understanding of the distinctive neural coding associated with visual stimuli, as they traverse this hierarchical landscape, remains elusive. In this study, we leverage the comprehensive Allen Visual Coding dataset and utilize the capabilities of deep learning neural network models to study the question of neural coding in response to dynamic natural visual scenes across an expansive array of brain regions. We find that our decoding model adeptly deciphers visual scenes from neural spiking patterns exhibited within each distinct brain area. A compelling observation arises from the comparative analysis of decoding performances, which manifests as a notable encoding proficiency within both the visual cortex and subcortical nuclei, in contrast to a relatively diminished encoding activity within hippocampal neurons. Strikingly, our results reveal a robust correlation between our decoding metrics and well-established anatomical and functional hierarchy indexes. These findings not only corroborate existing knowledge in visual coding using artificial visual stimuli but illuminate the functional role of these deeper brain regions using dynamic natural scenes. Consequently, our results proffer a novel perspective on the utility of decoding neural network models as a metric for quantifying the encoding of dynamic natural visual scenes, thereby advancing our comprehension of visual coding within the complex hierarchy of the brain.

## I. INTRODUCTION

Over the course of several decades, extensive research has yielded profound insights into the neural encoding of various attributes within artificial visual stimuli, encompassing features like the direction of moving gratings. This wealth of knowledge has shed light on the encoding mechanisms employed by visual neurons located across distinct regions of the brain, particularly within the early visual processing systems, which encompass the retina [1]–[5], the lateral geniculate nuclei (LGN) [6]–[10], and the primary visual cortex (V1) [11]–[14]. Nonetheless, a formidable challenge remains in elucidating how neurons distributed across various regions of the brain represent natural scenes, comprising natural images and videos [15]–[18]. This challenge is particularly pronounced when investigating the neural coding of dynamic natural scenes, such as videos [19]–[24]. The visual system stands as a quintessential neural domain, bearing paramount significance in mediating interactions with the external environment. A considerable expanse of the cerebral cortex is allocated to the processing of visual information [25]–[27]. This intricate cascade of information processing unfolds as we receive copious sensory inputs from the external world and engage in higher-order cognitive functions following the intricate orchestration of these inputs across various brain areas.

The retina serves as the inaugural site of the intricate conversion of diurnal visual scenes into electrical signals, marking the inaugural phase in the trajectory of visual coding. These visual signals traverse the brain in the form of neural spikes, commencing their journey with the retinal ganglion cells, progressing through the LGN located in the thalamus, and ultimately arriving at the visual cortex. Within the realm of the visual cortex, two discernible information pathways, namely the dorsal and ventral streams, have been suggested to come into play [28], [29]. In both streams, the initial point is the V1, serving as the pivotal site of early neural processing [30]. In the dorsal stream, where intricate spatial and locational information processing occurs, neural signals undergo further transmission to the anterior pretectal nuclei (APN) [31], [32]. Simultaneously, within the ventral stream, which contributes to memory formation, information takes a divergent route, eventually reaching hippocampus [33]. The elucidation of how visual signals are encoded and subsequently decoded within the cerebral confines is a paramount inquiry in the field of vision research [34]. In particular, the revelation of how visual scenes traverse a hierarchical neural structure within the brain constitutes a foundational inquiry that holds profound implications not only for the realm of vision but also for the broader computational principles governing the functioning of neurons and neuronal circuits [4], [19], [35].

Over the past several decades, the mechanism governing the encoding of visual scenes has undergone comprehensive investigation, culminating in a wealth of insightful perspectives on the encoding of specific visual attributes, including but not limited to luminance contrast, directional motion and velocity [4], [13], [35]. Moreover, these inquiries have extended to encompass more intricate and holistic visual scenes, such as natural images and videos [16], [19], [36]. In contrast to encoding, the challenge of decoding visual information from neural signals has been predominantly approached as an engineering perspective. In this context, an array of methodologies and models have been developed, primarily oriented toward addressing classification tasks related to object categorization [37]–[39], as well as the endeavor of reconstructing pixel-level images [40]–[50]. These recent studies in decoding and the reconstruction of pixel-level imagery provide fresh perspectives that hold significant implications for the evaluation of visual neuroprosthetic devices and the advancement of vision restoration [51]–[54].

Nevertheless, the interaction of encoding and decoding methodologies within the context of visual coding, especially pertaining to dynamic natural scenes, remains an area marked by limited exploration. In this study, we undertake the endeavor of uniting these facets to offer a comprehensive perspective on visual coding. To accomplish this, we investigate a well-established and robust resource, the Allen visual coding experimental dataset [55], utilizing a deep learning neural network decoding model [49]. Our exploration delves deeply into the Allen dataset, exploring it systematically to unravel the intricacies of visual coding distributed across a wide spectrum of hierarchical brain regions. These encompass three distinct regions within the nucleus, six segments within the visual cortex, and four divisions within the hippocampus. Harnessing the power of our decoding model, we embark on the intricate task of reconstructing every individual pixel within the images from the corresponding neural spikes. In this pursuit, we ascertain our capability to accurately decode video pixels from the neural activity of neurons located within the nucleus and the visual cortex, while discerning a significant decrease in decoding accuracy within the hippocampus. Notably, our findings reveal a strong correlation between our decoding accuracy and the classical encoding metrics obtained from experiments employing artificial stimuli. Furthermore, these findings extend to encompass the alignment of our decoding accuracy with the established anatomical and functional hierarchy indexes identified through experimental investigation. These significant outcomes underscore the substantive meaning and competence of our decoding model in deciphering visual information embedded within neural signals. Consequently, our study introduces an innovative approach to quantifying the extent to which visual information derived from dynamic scenes is encapsulated within neural signals from distinct cerebral regions.

## II. RESULT

### A. Encoding and decoding dynamic visual scenes in different brain areas

To elucidate the hierarchical organization of visual processing within the intricate pathways of the brain, we used a robust and expansive experimental dataset, the Allen Visual Coding dataset [55]. This dataset providesrecordings from thousands of neurons within mice, captured simultaneously via neuropixels, across a diverse array of brain regions (Fig. 1). Our investigation focused on three principal brain regions, encompassing a total of 13 identified brain areas. These regions are delineated as the visual cortex (comprising VISp, VISl, VISrl, VISal, VISpm, and VISam) located at the uppermost tier of the brain, the hippocampus (encompassing CA1, CA3, DG, and SUB) located at an intermediary level, and the nucleus (encompassing LGN, lateral posterior nucleus - LP, and APN) located in the deeper recesses of the brain (Fig. 1A).

**Fig. 1:**
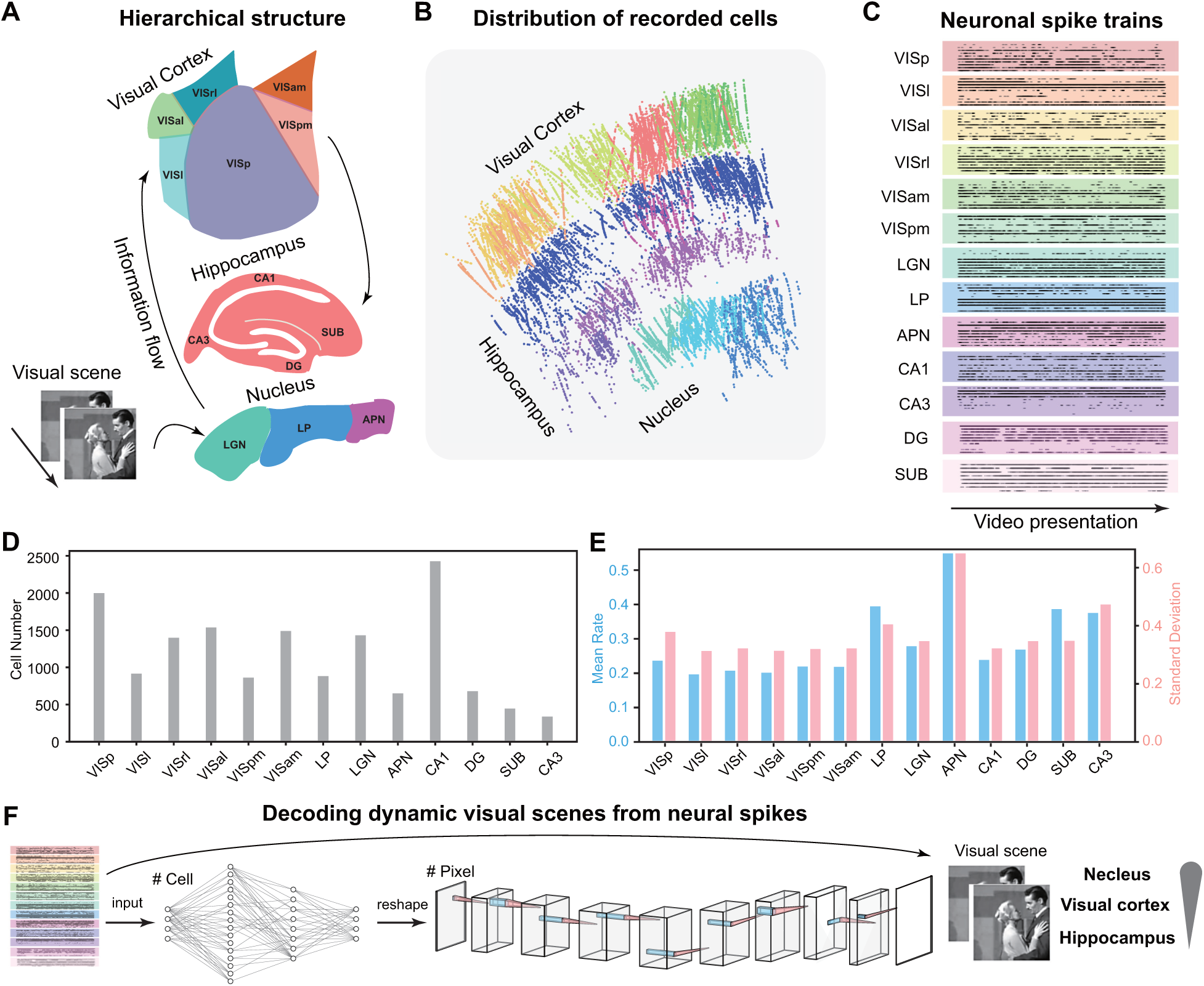
Encoding and decoding dynamic visual scenes using the Allen Visual Coding dataset. (A) A hierarchical structure of 13 brain areas in three brain regions of the visual cortex, hippocampus, and nucleus, was recorded in response to dynamic videos. The visual information flow is indicated by arrows. (B) The distribution of cell locations recorded in response to videos on multiple neuropixels. Each datapoint is an individual cell. (C) Example neuronal spike trains of 10 cells in each brain area in response to a video presentation. (D) The distribution of cell numbers in each brain area. (E) The firing activity showing means and standard deviations (SDs) of the spike counts averaged over the entire duration of video stimuli in each brain area. (F) The decoding workflow. A deep learning neural network decode takes the input of neural spikes and outputs images. The decoder performance indicates how much visual information is encoded by different brain areas.

Within the Allen dataset, a diverse pool of visual stimuli was deployed, ranging from well-designed artificial scenes, such as drifting gratings designed to assess fundamental tuning properties, to dynamic video scenes that have not been extensively investigated. Neurons were systematically recorded in response to these stimuli, with multiple neuropixels capturing a large set of cells distributed across a wide spectrum of brain areas (Fig. 1B). Notably, the spiking activity observed in response to the video presentations exhibited dynamic temporal patterns both within individual cells and across the cell population (Fig. 1C).

While previous studies have successfully characterized the encoding properties of artificial visual scenes, including the tuning of direction and orientation, across various brain regions [55], the encoding of dynamic video scenes remains comparatively less understood. Here we selected a subset of cells exhibiting firing activities in response to video stimuli. The cell count within this subset exhibited variation, ranging from 355 cells in CA3 to 2443 cells in CA1, with most brain regions comprising more than 800 cells (Fig. 1D, Table S1). The firing rates, encompassing both means and standard deviations of spikes, manifested a diverse range of values yet demonstrated consistency across the distinct brain areas (Fig. 1E, Fig. S1). This comprehensive dataset with high-quality neuronal recordings and dynamic video stimuli empowers our exploration into the representation of dynamic natural scenes across different brain regions.

The core focus of our study is the direct decoding of dynamic visual scenes from neural spike data. To achieve this, we leverage our previously developed deep learning model, which has been validated to decode image pixel data from neuronal spikes originating in the retina [49]. Here, we extend the application of this model to explore the intricacies of this expansive dataset. Our model, designed as an end-to-end deep learning neural network (Methods), receives sequences of spikes from a population of cells as input and generates, as output, the corresponding pixels of video frames associated with these spikes. By quantifying the fidelity and quality of the reconstructed images, we are equipped to study the extent to which visual information is encoded within distinct brain areas. It is a reasonable expectation that the nucleus and the visual cortex should exhibit a substantial information load, whereas the hippocampus is expected to bear relatively less visual information (Fig. 1F).

### B. Decoding model as a metric of visual scene encoding

Utilizing a consistent deep neural network framework, we embarked on the task of decoding the same visual images by inputting distinct neural spike data originating from various brain regions. Our model exhibited a superb capability for decoding and reconstructing image pixels with a high degree of precision (Fig. 2A). To evaluate the quality of the decoded images, we employed two widely accepted quantitative metrics: Structural Similarity (SSIM) and Peak Signal-to-Noise Ratio (PSNR). Both of these metrics were adeptly used as measures for assessing the performance of our decoding model, consistently yielding evaluations of the reconstructed image quality (Fig. 2B, Fig. S2).

**Fig. 2:**
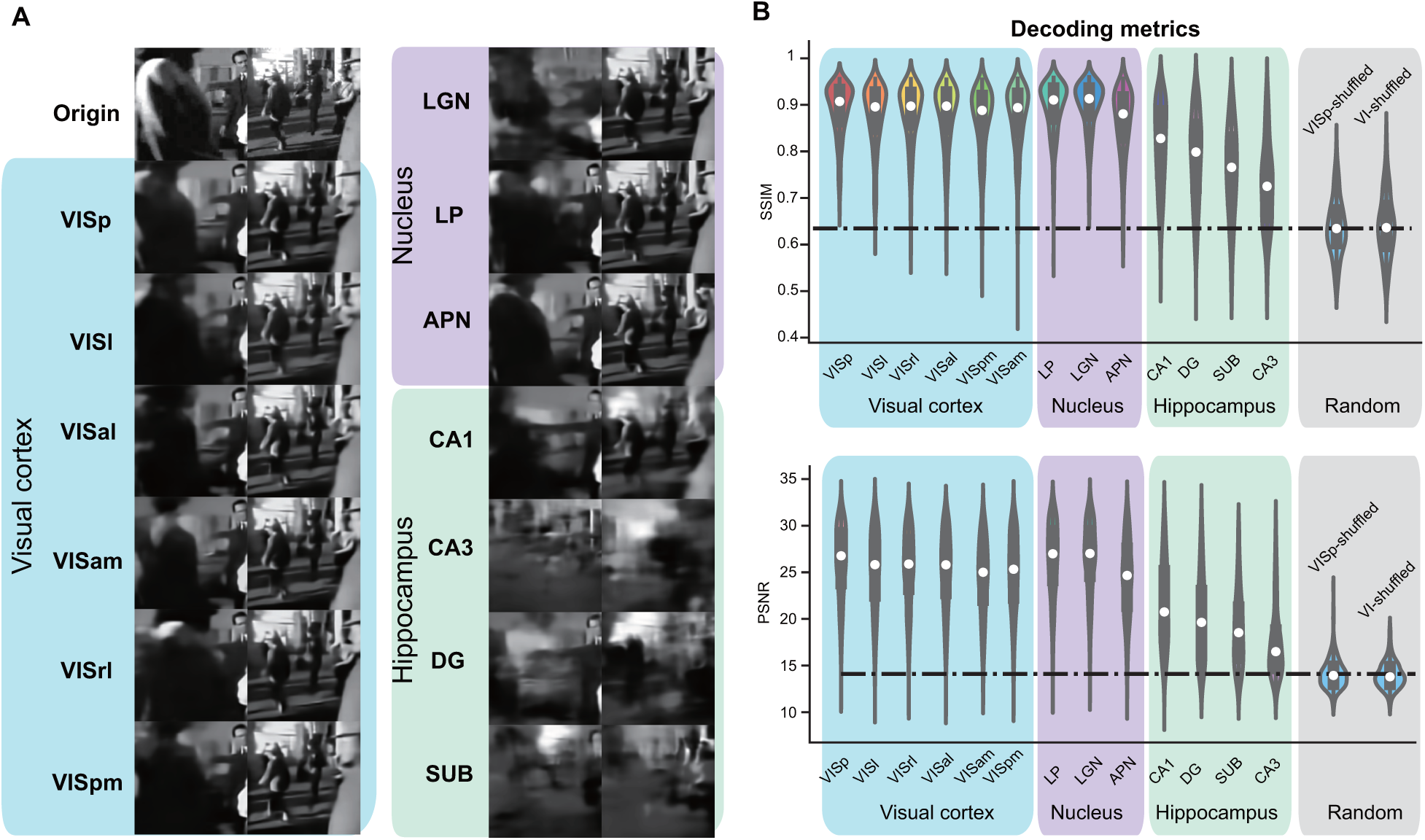
Decoding dynamic visual scenes in individual brain areas. (A) Example of decoded video frame images using spikes of each individual brain area. The original images are on the top (Origin). The decoded images from each brain area are colored according to the visual cortex, nucleus, and hippocampus. (B) Decoding metrics, SSIM and PSNR, indicate the quality of decoded images in different brain areas. The random cases serve as decoding baselines (dash lines), using two shuffling scenarios, shuffled spikes in the primary visual cortex (VISp-shuffled), and all six areas of the visual cortex (VI-shuffled). The values in violin plots are computed with 400 test images in this and the following figures.

A pronounced trend of visual coding decay was revealed as we traversed the hierarchical landscape of the visual information pathway. The primary visual cortex (VISp) emerged as the most proficient in rendering images with a high degree of fidelity, faithfully capturing the details of the stimuli. Furthermore, within the sub-regions, each of the six distinct brain areas located within the visual cortex, as well as the LGN and LP within the thalamus, and the APN within the midbrain, exhibited decoding results of considerable quality. These regions could effectively reconstruct a significant portion of the original image details. In contrast, results obtained from the hippocampus areas were notably inferior, reflected in diminished values of decoding metrics and a substantial loss of fine-grained details from the original stimuli.

To determine the significance of the decoding metrics obtained, we performed an experiment involving the random shuffling of spikes within the primary visual cortex (VISP-shuffled) and across all six regions of the visual cortex (VI-shuffled). This process effectively perturbed the temporal relationships between the spikes and the stimulus images, rendering the reconstruction process entirely random without any meaningful information. The decoding metrics SSIM and PSNR were measured for the data resulting from the shuffled spikes, serving as a baseline for comparison. Our decoding results consistently outperformed the shuffled baseline, underscoring the meaningful nature of our decoding approach and its capacity to faithfully reflect the extent of visual information encoded within neural spikes (Fig. 2B).

### C. Hippocampus encodes less visual information

To undertake a comprehensive quantification of how visual scenes are encoded across various brain areas, we aggregated all nearby sub-areas into three cohesive and interconnected macroscopic brain regions: the visual cortex, nucleus, and hippocampus. Using the collective neuronal spikes originating from these three distinct brain regions, we conducted a thorough examination of decoding outcomes and subsequently compared their performance across these regions (Fig. 3). In particular, the example decoded images in each region show a striking level of precision in image reconstruction, distinguishing them significantly from the results of shuffled spikes (Fig. 3A). These favorable results were further validated by the decoding metrics SSIM and PSNR, which underscored the superior performance of both the visual cortex and the nucleus in contrast to the hippocampus (Fig. 3B).

**Fig. 3:**
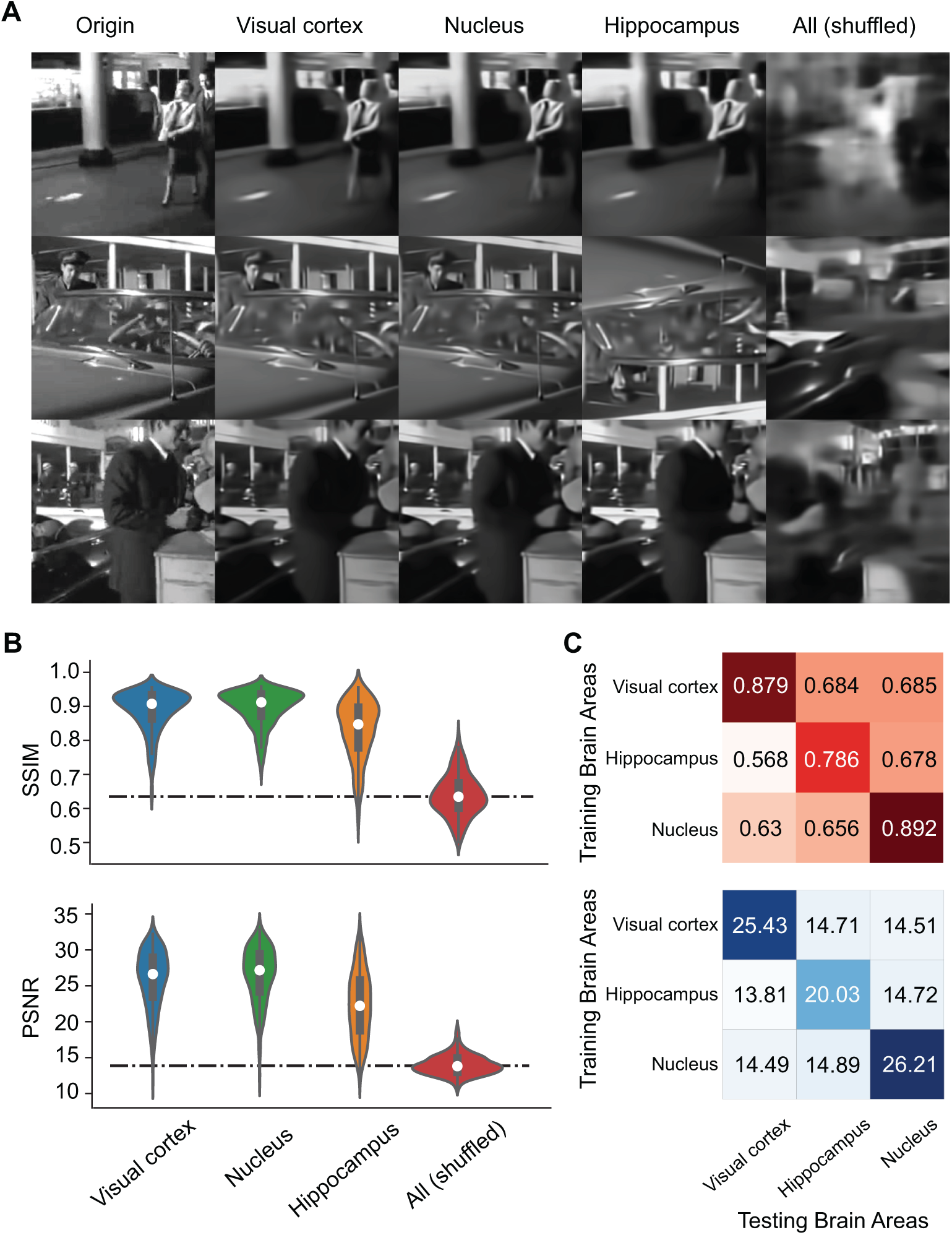
Decoding results in each brain region. (A) Samples of original images (Origin) and decoded images in each combined brain region (Visual cortex, nucleus, hippocampus). The decoded results in shuffled spikes across all brain regions listed as a baseline. (B) Corresponding decoding metrics in each case of (A). SSIM and PSNR metrics in each brain region and baseline with shuffled data. (C) Decoding matrices where the diagonal elements are the values in (B) and the off-diagonal elements are the values of model generalizability, e.g., using the models trained on each brain area to predict other test brain areas. Marked values are the means of over 400 test images in this the following figures.

To explore the generalizability of our decoding models, we applied a model trained on one specific brain region to data originating from different regions. This approach facilitated the construction of a decoding matrix (Fig. 3C), wherein the diagonal elements reflect the model’s performance when trained and tested with data from the same brain region. In contrast, the off-diagonal elements represent the model’s generalizability, where the performance was trained with one region’s data and tested on another. Notably, we observed that the decoding models exhibited a remarkable degree of specificity, with relatively low generalizability. The decoding metrics associated with off-diagonal elements closely match the performance observed in the shuffled baseline, underscoring the distinctive nature of our decoding models (Fig. 3C).

### D. Robust decoding with a small set of cells

In pursuit of a more detailed comparison between different brain areas, we systematically replicated our decoding procedure, by employing controlled cell numbers from each region. To ensure a limited sampling representation, we randomly selected 800 cells from all six areas within the visual cortex, as well as the LGN, LP and CA1. This selection was made with the understanding that other regions possessed fewer than 800 cells. Remarkably, the decoding results derived from this subset of 800 cells closely match those obtained with the full number of cells (Fig. 4A). This observation was further substantiated by the decoding metrics SSIM and PSNR. Furthermore, we explored two distinct scenarios wherein we mixed areas from the visual cortex, with or without VISp. In each case, decoding with 800 cells consistently yielded akin results. However, the decoding performance with CA1 remained notably inferior when contrasted with that of the visual cortex, LGN, and LP. The decoding models trained on individual brain areas exhibited remarkable specificity, demonstrating inferior performance when transferred to test data from other brain regions (evident in the off-diagonal metrics, which closely mirrored the shuffled baseline, as illustrated in Fig. 4B).

**Fig. 4:**
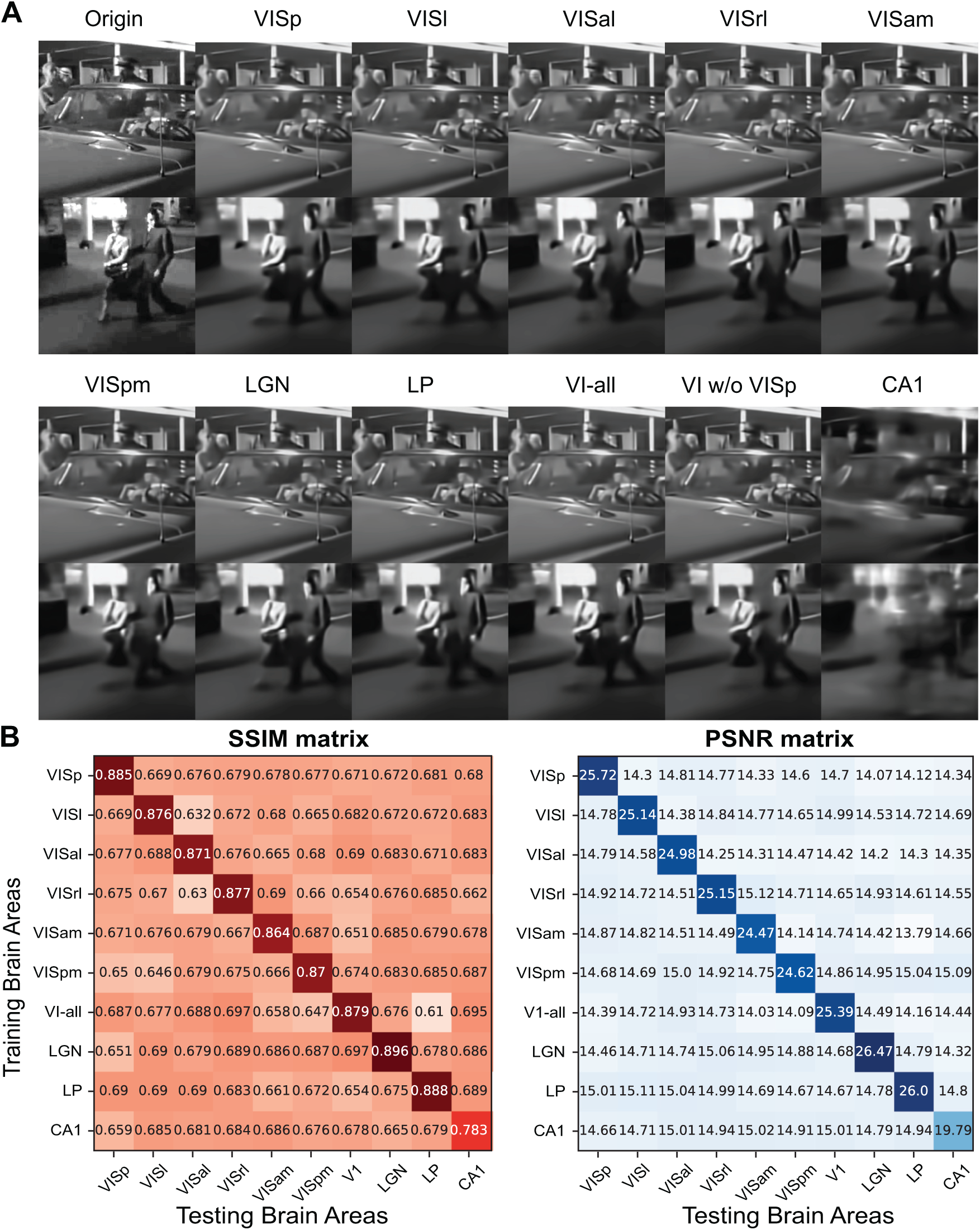
Decoding results are consistent with the same number of cells. (A) Examples of decoded images with 800 cells of each brain area and two combined areas (VI-all: all combined six areas in the visual cortex; VI w/o VISp: combined five areas of the visual cortex without VISp). (B) Decoding metrics of SSIM and PSNR. Matrices show the decoding values using models trained on each brain area while testing on the same (diagonal) and different (off-diagonal) areas.

We then investigate the influence of cell quantity on the decoding outcomes. Cells were randomly selected in varying numbers, ranging from 50 to 2000 within VISp, and the decoder was trained using differing numbers of cells under the same scheme (Fig. 5A). Not surprisingly, the quality of decoding decreased when fewer cells were employed. This examination was extended to encompass all brain regions, and it became evident that the decoding metrics exhibited a consistent upward trend with an increase in the number of cells included for model training. However, an intriguing phenomenon was observed - the performance reached a state of saturation when there are enough number of cells incorporated (Fig. 5B). Specifically, the visual cortex and nucleus regions appeared to reach a saturation point with approximately 500 cells, whereas the hippocampus necessitated around 1000 cells for the same effect. This observation implies that cells within the nucleus and visual cortex exhibit a greater degree of specialization in processing visual information in contrast to those within the hippocampus.

**Fig. 5:**
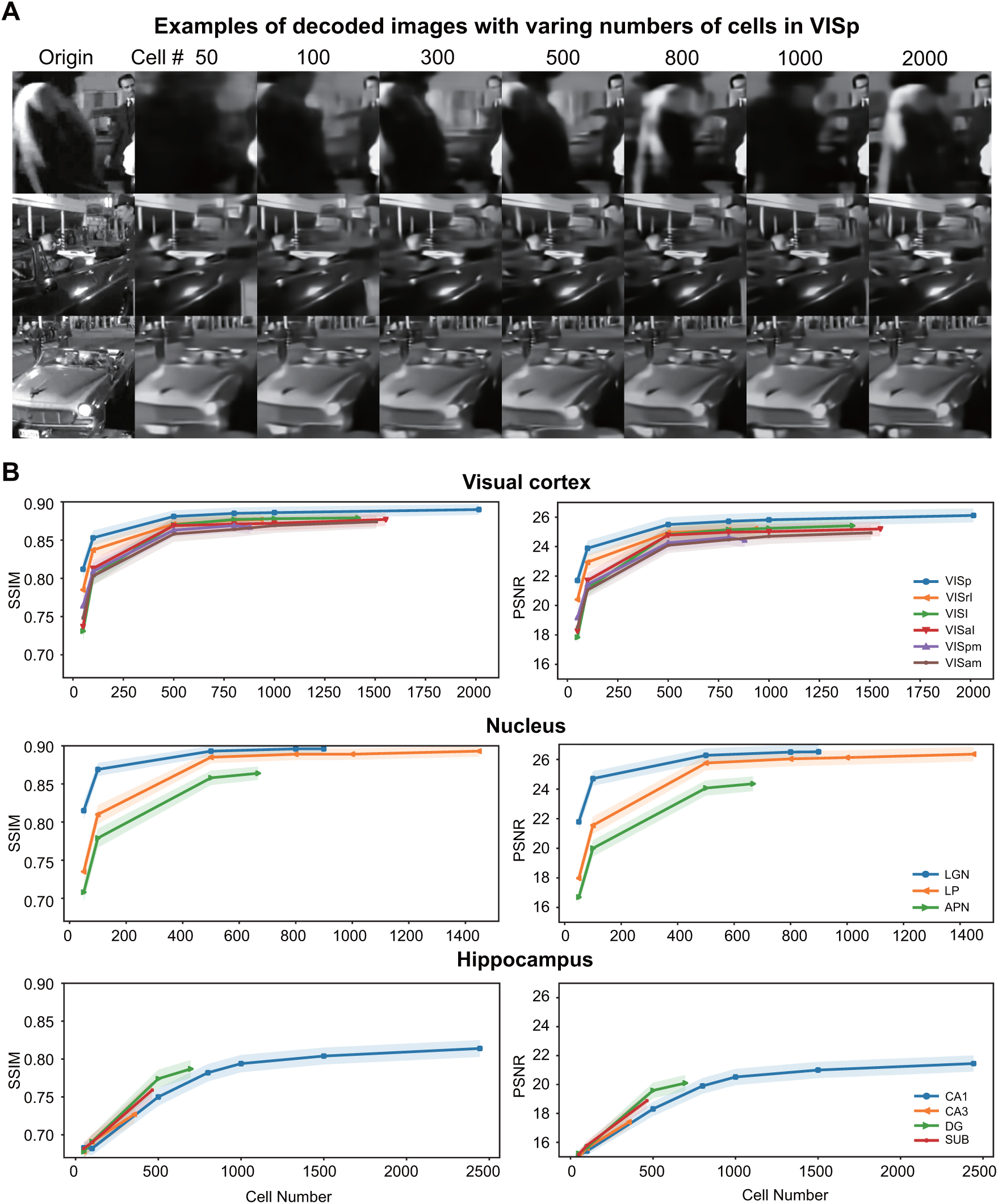
Decoding results are saturated with a small set of cells. (A) Examples of decoding images with different numbers of cells (50-2000) in VISp. (B) Decoding metrics (mean*±*SD) are convergent over an increasing number of cells in each brain area.

A notable observation was that brain areas that exhibit higher decoding performance consistently outperformed other regions (Fig. 5B). The performance curve of VISp consistently ranked highest among all segments of the visual cortex, even with a limited number of cells for decoding. Similarly, the LGN consistently exhibited superior decoding results compared to LP and APN. In contrast, the hippocampus displayed a lower decoding performance, even when a substantial number of CA1 cells were utilized. Even though the Allen dataset featured fewer cells in CA3, DG, and SUB, it was anticipated that the decoding performance of these hippocampal regions, with a greater number of cells, would remain suboptimal, comparable to that in CA1. These findings suggest that an accurate decoding of visual scenes can be achieved with a relatively small number of cells, depending on the brain areas. In terms of image pixel decoding, the information contained in the spike data of each brain area appears to possess a certain degree of redundancy. The total information derived from the original stimuli, funneled through each brain area, seems to be a constant quantity. Even when a substantial number of cells were employed in CA1, the volume of information it encompassed was not on comparable to that decoded from a limited number of cells in V1. Collectively, our results suggest that cell quantity is not a deterministic factor in the decodability of visual scenes.

### E. Decoding metrics are consistent with the encoding indexes of artificial scenes

The Allen dataset encompasses neural activity responses to a diverse array of stimuli, including static and drifting gratings, serving as a valuable resource for calculating neural selectivity towards different orientations and directions. To gain a deeper comprehension of how decoding performance may elucidate encoding capabilities, we characterize the relationship between de-coding performance and cell selectivity tuning. We employed three classical indexes widely used in the field of visual coding (Fig. 6): orientation selectivity to static grating stimuli, orientation selectivity to drifting gratings, and directional selectivity to drifting gratings.

**Fig. 6:**
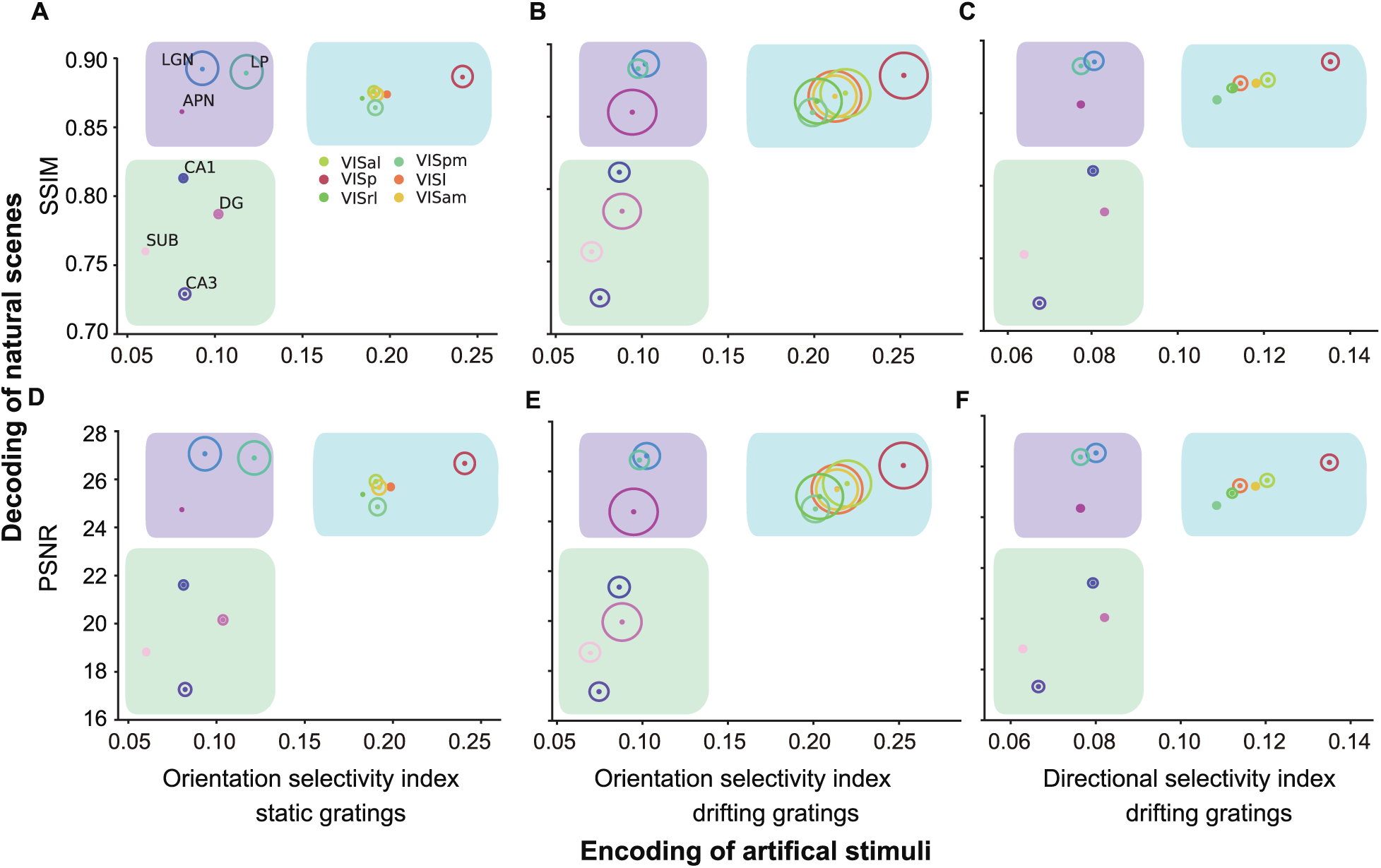
Decoding of natural scenes is consistent with the encoding of directional featured-based artificial stimuli. The relationship between natural scene neural activity image reconstruction performance and directional visual feature cell selectivity indexes. Natural scene decoding performance metrics SSIM (top; A-C) and PSNR (bottom; D-F) are plotted against (A) orientation selectivity indexes to static gratings, (B) orientation selectivity indexes to drifting gratings, and (C) directional selectivity indexes to drifting gratings. Solid datapoints are means. The circle size is proportional to the SD of different selectivity indexes.

Within the visual cortex, neurons demonstrated a pronounced tendency for high orientation and directional selectivity tuning, a propensity that is significantly more pronounced compared to neurons within the LGN, LP and APN, consistent with established conventions in the field [7], [13]. In contrast, all hippocampal areas exhibited both low decoding metrics and diminished orientation selectivity. This diversity generates a distinct pattern, categorizing regions into three distinct clusters: the visual cortex, marked by its high orientation encoding capabilities coupled with a strong decoding performance; the nucleus, marked by low encoding but a strong decoding performance; and the hippocampus, marked by both low encoding and decoding performance. Consequently, these findings indicate the striking consistency in the quantitative relationships between our decoding metrics derived from natural scenes and the encoding tuning properties traditionally associated with artificial scenes. Therefore, our decoding model serves as a functional metric for the study of the encoding of natural visual scenes.

### F. Decoding of natural scenes across the visual hierarchy

Recent investigations have revealed the presence of a hierarchical organization within the visual cortex, established through an analysis of regional connections [30], [55] (Fig. II-F A). To delve deeper into the hierarchy of natural scene decoding, we conducted an in-depth examination of the regions within the visual cortex (Fig. 6). This inspection revealed a substantial positive correlation between the SSIM and PSNR decoding metrics and the selectivity indexes associated with artificial stimuli in the six areas of the visual cortex (Fig. II-FB-D). Notably, a stronger correlation emerged between decoding performance and cell selectivity for drifting gratings compared to static gratings (Fig. II-FB-D), signifying a more prominent role for the visual cortex in encoding orientation and direction for dynamic vision compared to static vision.

**Fig. 7:**
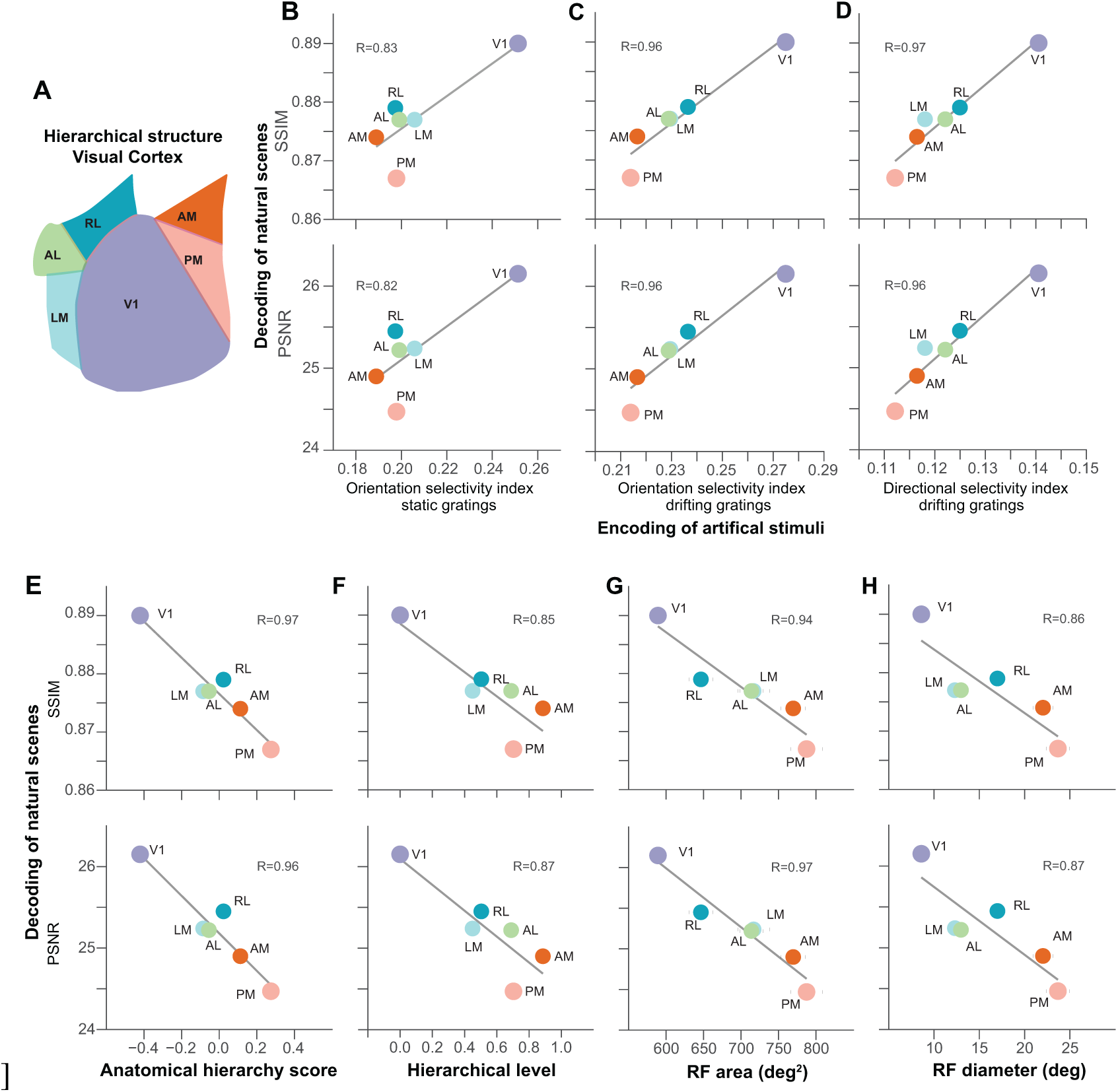
Decoding of natural scenes in the visual cortex hierarchy. (A) Diagram of the visual cortex.(A) Diagram of mouse visual cortex, showing the anatomical layout of the regions. Regions are synonymous with previous analyses: V1 (VISp), LM (VISl), RL (VISrl), AL (VISal), PM (VISpm), AM (VISam). (B-D) The relationship within the visual cortex between decoding metrics SSIM (top) and PSNR (bottom), and directional visual feature cell selectivity indexes (B) orientation selectivity using static gratings, (C) orientation selectivity to drifting gratings, and (D) directional selectivity to drifting gratings. (E-H) Correlation between decoding performance metrics SSIM (top) and PSNR (bottom) with (E) anatomical hierarchy score [56]; (F) hierarchical level [30]; (G) receptive field (RF) area [55]; and (H) RF diameter [30]. Data are presented as mean values. R is the Pearson correlation coefficient. For all correlations P*<*0.05.

Subsequently, we further dissected the correlation between the decoding metrics and the previously established hierarchical structure within the visual cortex [30], [55], [56] (Fig. II-F E-H). The decoding metrics exhibited a high correlation with the anatomical hierarchy score [56] (Fig. II-F E). Similarly, the hierarchical level values obtained from another independent anatomical tracing study, which employed electrophysiological methods, displayed a robust correlation with both decoding metrics [30] (Fig. II-F F). Furthermore, we analyzed another aspect of visual neuron properties, receptive field (RF) size, which has been previously demonstrated to expand in higher-order areas as visual features are aggregated [30], [55]. Intriguingly, we observed a highly significant correlation between decoding metrics and RF sizes, a correlation that was consistent across experiments conducted both by [30], [55] (Fig. II-F G-H). In particular, the correlation was more pronounced in the case of the Allen dataset (Fig. II-F E and G, R*>* 0.9) compared to the data presented by [30] (Fig. II-F F and H, R*<*0.9).

## III. Discussion

Our primary focus revolves around the direct decoding of dynamic visual scenes from neural spikes, employing an end-to-end deep learning model as the decoding mechanism. Leveraging the valuable resource provided by the Allen dataset, we systematically explored how visual scenes find their neural representation within diverse brain regions. Specifically, we endeavor to address the longstanding question concerning the extent to which neurons in the hippocampus encode visual information. Traditionally, it has been a well-established notion that the primary contributors to visual information encoding are the visual cortex and the LGN, while the hippocampus, in its capacity for information integration towards associative learning and memory formation, encodes a fraction of visual information. However, our novel decoding approach introduces, for the first time, a quantitative metric that allows for the intricate quantification of this established paradigm.

### A. Decoding visual scenes at the pixel level

Conventional paradigms in neuroscience, particularly within the domain of decoding studies, have traditionally focused on using neural signals to retrieve and classify information related to stimuli. These efforts have yielded valuable insights into the neural code [55], [57]–[59]. In recent years, a significant shift has been observed, with an emerging emphasis on the reconstruction of input images or videos using a diverse array of neural signals. These signals encompass functional magnetic resonance imaging (fMRI) data [38], [41], [42], [60], calcium imaging signals [48], [50], and neural spikes [3], [43], [45]–[47], [61]–[63]. In the present study, we break new ground by undertaking the task of decoding and reconstructing visual scenes across multiple brain regions, ranging from the nucleus and visual cortex to the hippocampus, with a specific focus on unraveling the intricacies of the visual hierarchy.

The fundamental question of how neurons in the brain encode visual signals stands as a cornerstone in the field of neuroscience. The prevailing notion entails that visual signals undergo initial encoding within retinal ganglion cells, further progressing through encoding and decoding processes in the LGN and the visual cortex. As these signals traverse deeper into the brain, they are subjected to processing that extracts more abstract, meaningful information for the purpose of learning and memory formation. Previously, our work demonstrated the capacity of deep learning models to decode image pixels from neural spikes in the retina [49]. In the current investigation, we apply a similar approach to address decoding using neurons in later stages of the visual pathway.

Of particular note, the LGN and LP, located at the forefront of the visual pathway following the retina, exhibit remarkable accuracy in decoding natural scenes. Their proximity to the initial stage of visual processing may account for their ability to retain a substantial amount of visual information. The six regions within the visual cortex and the APN in the midbrain also yield commendable decoding results. Given the crucial roles played by the primary visual cortex in both the dorsal and ventral visual streams and the involvement of APN in the processing of spatial location information within the ventral stream, it is reasonable to anticipate their need for a wealth of original stimulus information.

In contrast, the decoding results for the hippocampal regions, including CA1, CA3, DG, and SUB, appear notably less robust. This discrepancy could be attributed, in part, to their position at the far end of the visual pathway. It is conceivable that the processes of learning and memory formation do not necessitate an exhaustive retention of intricate pixel-level details from the original stimuli. Our findings align with the prevailing understanding that visual information undergoes hierarchical processing across distinct neural regions, each contributing to a unique facet of the visual information pathway.

The redundancy inherent in the representation of pixel-level information within each distinct brain area becomes apparent in our findings. Notably, we have observed that the use of 800 cells is sufficient to approximate decoding results that align closely with those obtained using the entire population of cells within a given brain area. Intriguingly, even when working with a sparse subset of cells, such as 50 or 100, our decoding outcomes remain reasonable and surpass those generated by random baseline measures. Furthermore, increasing the number of cells serves to reduce blurriness and enhance the overall quality of image details in our decoding results. It becomes evident that, once the cell count reaches approximately 800, or even as few as 500, the decoding outcomes are in line with those derived from a larger dataset of 2000 cells. However, when progressing into the hippocampus, we find that a more substantial number of cells, approximately 1000 to 1500 in the case of CA1, is requisite to attain a sufficiently reasonable level of pixel decoding. These empirical insights provide compelling evidence of the redundancy in the representation of pixel-level stimuli within each unique brain area. It is remarkable to note that even a sparse population of cells has the capacity to capture and convey the essential details of image pixels, underscoring the efficiency and robustness of neural encoding.

### B. Decoding visual scenes in hierarchical neural pathways

Information processing within the visual pathway unfolds along a hierarchical cascade involving sequential stages encompassing the retina, subcortical regions, and cortical areas. Within this intricate neural pathway, visual information undergoes a transformation and is distributed extensively to various regions of the brain. This cascade is widely believed to follow a hierarchical organizational principle, with higher-order brain regions performing more sophisticated computations involving increasingly encoded information [56]. A rich body of previous research, utilizing advanced techniques such as 2-photon calcium imaging [56] and electorphysiology [30], has established the presence of hierarchical organization within distinct regions of the mouse visual cortex. The visual hierarchy becomes notably manifest in the response latency as one traverses along the hierarchy, with higher-order regions exhibiting slower response times, thus corroborating previous findings derived from imaging studies [30], [56].

To unveil the capacity of decoding metrics in elucidating the information representation across this hierarchical visual pathway, we conducted a rigorous examination of their interplay with the functional anatomical structure. Our exploration involved three well-established encoding indexes, designed to capture the selective activity of neurons within various brain regions when exposed to artificial stimuli in the form of static and drifting grating patterns, as sourced from the Allen dataset. These indexes offer insights into how individual neurons are finely tuned to distinct visual features, such as orientation and direction [64]. Subsequently, the tuning properties of neurons were thoughtfully correlated to decoding performance metrics obtained through the presentation of natural visual scenes. This critical analysis aimed to uncover the intricate relationship between neural encoding properties, such as selectivity tuning, and the representation of real-world visual stimuli by neural decoding (Fig. 6).

Areas within the visual cortex and the hippocampus reveal a noteworthy correspondence between the reconstruction performance metrics SSIM and PSNR obtained from the decoding model and the selectivity of cells to the orientation and direction. These findings highlight that decoding performance metrics, as acquired from the presentation of natural scenes, are quantitatively linked to the tuning properties of cells for encoding orientation and direction with conventional artificial stimuli. Consequently, the decoding model offers a valuable metric for the comprehensive study of orientation-based encoding in natural visual scenes, transcending the realm of artificial stimuli. In contrast, the regions located within the nucleus exhibit high reconstruction performance while displaying low cell selectivity indexes. This observation is in harmony with prior research indicating a lower direction selectivity index in the LGN compared to V1 [7], [11], [64].

Nevertheless, it is imperative to emphasize that the collective cell selectivity within a region does not necessarily correspond to the overall amount of visual information retained. The mouse LGN, for example, receives input from various types of retinal ganglion cells, encompassing information that spans a wide spectrum of visual features [8]. Furthermore, neurons within the LGN of rodents and other mammals have been observed to manifest circularly symmetric receptive fields, in contrast to the elongated receptive fields found in V1 [7], [11], [64], [65]. While LGN neurons may indeed carry the necessary information to interpret orientation and direction, their selectivity appears to become more pronounced when their receptive fields converge within V1. Additionally, neurons in the rodent and mammalian LGN demonstrate marked selectivity to light intensity and linear spatial summation, in addition to their orientation and direction information [66], [67]. The human LGN, characterized by a layered structure, carries a highly diverse range of visual information emanating from both the parvocellular and magnocellular visual pathways, located in dorsal and ventral layers respectively [68]. Consequently, certain groups of cells within the region may exhibit orientation selectivity, while others are attuned to different facets of visual information, thus impacting the overall orientation and direction selectivity indexes.

In contrast, the visual cortex prominently exhibits an elevated degree of orientation and direction selectivity, implying the primacy of this form of information representation within the region. However, each distinct region within the visual cortex demonstrates varying selectivity indexes, with V1 emerging as the most selective for orientation and direction among artificial stimuli in the majority of cases. This aligns cohesively with the previously outlined functional hierarchy of the visual cortex, wherein V1, marked by its highest orientation selectivity, serves as the point of origin for an expansive network of orientation-selective cells originating from LGN inputs [7], [11], [64]. Subsequent regions within the visual cortex hierarchy exhibit weaker preferences for orientation, indicating their role in processing alternative forms of visual information and the transformation of visual data from V1 into more intricate representations. For instance, V3 has been suggested to demonstrate a greater preference for texture and pattern information [69]. Hence, the inclusion of additional selectivity metrics, such as those focused on texture selectivity or those unrelated to orientation and direction, would likely reveal distinctive trends among various regions.

An essential inquiry concerns the comparative reconstruction performance of the LGN and LP models when contrasted with V1. An equivalent performance may indicate that the LGN does not utilize other information besides orientation for enhancing reconstruction accuracy. In contrast, better performance could suggest the utilization of alternative visual information sources to enhance reconstruction performance. Moreover, higher reconstruction performance may imply the model’s capacity to employ spikes derived from alternative forms of visual information to augment reconstruction. While the current decoding model serves as a functional metric for the investigation of natural vision encoding beyond artificial stimuli, its competence in deciphering visual features other than orientation and direction remains enigmatic. Consequently, the examination of regions primarily relying on forms of visual information other than orientation, such as the visual cortex, may necessitate the application of alternative cell tuning metrics or decoding model architectures for the exploration and interpretation of the presence of diverse forms of visual information. Further studies are needed to illustrate these intriguing questions in more detail.

### Visual coding using deep learning models

Deep neural network (DNN) models have emerged as valuable tools in neuroscience research [70], particularly for visual coding using neuronal data from spikes to calcium imaging and fMRI [34], [50], [54], [71]–[74]. Seminal studies have demonstrated that DNNs can identify neuronal representations of visual features encoded in the visual cortex and inferior temporal (IT) cortex of primates [34], [71], [72]. In the mouse visual system, DNNs have aligned visual cortical areas with corresponding model layers using the same Allen Visual Coding dataset while focusing on neuronal responses to natural images [75]. Further applications of DNNs on additional data from the mouse visual system have provided deeper insights into visual coding [24], [76], [77]. While DNNs were initially designed to model the ventral visual processing pathway [34], questions remain regarding the adequacy of these models in fully explaining the underlying biological visual processes [78].

Similar to primates, mice can perceptually detect higher-order statistical dependencies in texture images, distinguishing between different texture families across visual areas in alignment with DNN predictions [79]. This observation implies that mouse visual cortex areas may represent semantic features of learned visual categories [80]. Beyond visual coding, the rodent hippocampus is suggested to have similar roles in learning and memory as observed in primates [81], [82]. In our study, we demonstrate that the same DNN reveals less pixel information in hippocampal neurons compared to the thalamus and visual cortex. This finding aligns with the general belief that the hippocampus encodes more abstract information, such as concepts [83]. Consequently, a pertinent follow-up question is how to decode this abstract information from mouse hippocampal neurons. Recent studies have shown that DNNs can decode semantic information from human fMRI data using advanced generative models. These models process latent embeddings and resample learned semantic information to generate or reconstruct new images with similar concepts [84]–[87]. Such generative models might be valuable for decoding semantic information represented by mouse hippocampal neurons. Additionally, developing models constrained by neuroscience knowledge could enhance decoding accuracy, offering new insights into the fundamental workings of the biological brain [88], [89].

Our current findings demonstrate that the decoding accuracy of natural scenes is closely correlated with encoding metrics derived from artificial scenes. While the interaction between encoding and decoding has been explored for simple stimuli such as directional selectivity [55], the investigation of complex natural scenes has been limited due to constraints in computational models and methodologies [22], [24]. Previous studies have shown that DNNs can be valuable tools for decoding and reconstructing pixel-level information of natural scenes [43], [44], [47], [50], yet these studies lacked a clear correlation with established encoding metrics. Our present work aims to elucidate this tight correlation, enabling the use of DNNs to quantify neuronal response patterns and compute the differences between patterns in response to both artificial and natural stimuli.

Our approach extends conventional metrics for assessing the distance between spike trains [90], [91], incorporating the capability to handle natural scenes. Consequently, our model provides practical and quantitative metrics for characterizing the quality of vision restoration based on neuronal responses treated by neuroprostheses [51], [52]. This advancement offers a robust framework for evaluating and enhancing neuroprosthetic treatments.

### Limitations

In the current work, our DNNs are based on convolutional neural networks that process video images frame by frame, without incorporating temporal information. In neuroscience, neuronal dynamics are characterized by complex temporal patterns. Including temporal information could potentially yield deeper insights. Our initial aim was to utilize the temporal sequence of visual processing to develop a spatiotemporal model using video based on continuous frames. However, recent advances in deep learning models for computer vision indicate that image-based foundation models consistently outperform video-based models on most video understanding tasks [92]. Consequently, models developed by analyzing static images can effectively address dynamic video tasks in a frame-by-frame manner, without the need for temporal information [93]. Consistent with these observations, we found that our current approach of analyzing video frame-by-frame yields better decoding results. Although our current model does not fully utilize the temporal sequence information, it provides a practical method for studying dynamic visual scenes and neuronal responses. Nonetheless, decoding dynamic continuous scenes in the brain while considering temporal information could offer more profound insights. Future work is needed to investigate temporal dynamics comprehensively.

It is well known that temporal delays exist across brain hierarchical areas in processing visual information [55]. For instance, deeper areas like the hippocampus respond to visual stimuli with a delay of several tens of milliseconds. To assess the impact of these delays on decoding capabilities, we reconstructed images using frames preceding the neuronal response and observed minimal effects (Fig. S3). However, our current model may not be suitable for examining how delays influence the processing and representation of visual information. At this stage, our decoding metrics reflect the hierarchical delays between brain areas, with reduced reconstruction errors being more pronounced in deeper areas. Future work will need to develop alternative modeling approaches that can investigate the interpretability of decoding performance while accounting for temporal delays.

Neuron dynamics across all areas of the mouse visual system can be modulated by behavioral variables and other contextual signals [24], [94]–[97]. The experimental settings of the Allen data involved anesthetized mice, thereby minimizing the influence of contextual differences. Future studies should consider how visual scenes are represented in mice located in more natural scenarios, with additional variables taken into account.

## Acknowledgment

We would like to thank Zhile Yang for the helpful discussions. This work was supported by the National Natural Science Foundation of China 62176003 and 62088102 and Beijing Nova Program 20230484362 (ZY) and the MOST of China 2022ZD0208604 and 2022ZD0208605 and National Natural Science Foundation of China Grants T2325008 and 820712002 (JZ), and Royal Society Newton Advanced Fellowship of UK Grant NAF-R1-191082 (JKL).

## Data Availability

The data used in this work are publicly available recordings from the Visual Coding - Neuropixels dataset, provided by the Allen Institute for Brain Science.

The stimuli images and neural data are available at:

https://portal.brain-map.org/circuits-behavior/visual-coding-neuropixels.

The code used to generate the results in this paper is available at https://github.com/beizai/Decoding.

## IV. Methods

### A. Experimental dataset

The extracellular electrophysiology dataset we use is from Allen Visual Coding [55]. This data set has 32 sessions of experiments. Each session contains three hours of total experiment data with the same procedure in different mice. Three hours of data include spiking responses to a battery of different stimulus scenes, including gratings and two movie clips whose lengths are 30 and 120 seconds. In this study, we used the 120-second movie with a total of 3600 frames, where each frame image has a size of 304*608 pixels. We counted those cells that release spikes under the movie clips and calculated the average number of spikes each movie frame generated in all sessions, in order to avoid the influence that every session contains different brain areas and every brain area contains different cell units in different sessions. If one cell does not release spikes in a session, then it is not included in our analysis. In the 32 experiments sessions, there are 25 brain areas containing more than 20000 cell units. Among them, we selected the brain areas with more than 300 cells, and the numbers of cells and their corresponding brain areas are shown in Table S1.

### B. Decoding model

The decoding model used in this study is similar to our previous models [49]. The first part of the network is a multilayer perceptron (MLP) with four layers of full connection layers. The size of the first layer is the number of input cells. Spike data of each cell corresponds to one neural cell in the network. The middle two layers are hidden layers, the size of which is 16384 and 8092, using the ReLU function as an activation function. The size of the final layer is 4096, outputting a 64*64 intermediate image.

The intermediate images from the MLP pass through a typical convolutional neural network and then get the final decoding result. The convolutional neural network is divided into two parts. The first part contains four convolutional layers, using convolution and down-sampling to reduce the image size. The sizes of four convolution kernels are (256, 7, 7), (512, 5, 5), (1024, 3, 3), and (1024, 3, 3). Stride sizes are all (2, 2). After this part, the network will reserve the main information from the input intermediate image and filter redundant noise. The second part contains six convolutional layers and six up-sampling layers. The convolution kernel sizes are (1024, 3, 3), (512, 3, 3), (256, 3, 3), (256, 3, 3), (128, 5, 5), (1, 7, 7). The stride sizes are all (1, 1). Sizes of up-sampling layers are all (2, 2). After the second part, we can decode and reconstruct the features of the original stimuli and get a reconstruction image with 256*256 pixels. Between every two convolutional layers, there are batch normalization, ReLU activation function, and dropout layer with 0.25 probability.

All decoding experiments used the same settings. Due to the high resolution of the original natural movie, we processed the movie by cropping to select the middle part of the images, such that the images were reduced to a resolution ratio of 256*256 pixels for our decoding model. With all 3600 image frames, we broke down the temporal order and randomly selected 3200 frames as the training and validation set and 400 frames as the test set. The loss function of the network is the mean squared error (MSE) between the reconstructed image and the original image. The training uses Adam optimizer with a learning rate of 0.001. The batch size is 16 and the epoch number is 400. All experiments are conducted on a Nvidia RTX 3080. The total parameter number is about 300 million and a single training costs about 200 minutes.

### C. Decoding metrics of natural scenes

We used two popular quantitative metrics to compare the decoded images with the original stimulus images of natural scenes: structural similarity (SSIM) and peak signal-to-noise ratio (PSNR). SSIM calculated the difference between the luminance, the contrast, and the structure between the two images. It is based on the hypothesis that human vision feels distortions by extracting the structure information from the images, whose value is between -1 and 1, proportional to the quality of the image. PSNR measures the degree of image distortion by dividing the maximum difference by the MSE between pixels from the reconstructed images and the original images. PSNR is an absolute value such that the bigger it is, the less distortion the result has. All the violin plots use the decoding metric values over 400 test images. The single values of the decoding metrics in all the other figures are the mean metric values over 400 test images.

### D. Encoding indexes of artificial scenes

We used three cell selectivity tuning indexes established in visual coding: orientation selectivity to static grating stimuli (OSI SG), orientation selectivity to drifting gratings (OSI DG), and (C) directional selectivity to drifting gratings (DSI DG). All these three indexes were measured and provided in the Allen data and the details of each index and experimental protocols were provided in [55]. We calculated the means of each index with all the cells that are in response to these grating stimuli. The means were then used as an overall measurement of cell tuning selectivity in different brain areas (mean values are listed in Table S2) to compare our decoding metrics of dynamic videos.

### E. Hierarchy indexes of visual cortex

Two types of hierarchical indexes of six areas of the visual cortex were used in this study. For fair comparison and validation between different experimental results, we used both types of hierarchy that were provided by recent studies of the Allen data [55], [56] and a separate study [30]. The first is the anatomical hierarchy index. The Allen data provides the anatomical hierarchy score of a large number of brain areas [56]. We also used another index named hierarchy level that is measured in a recent separate study [30]. Both measures have been shown to be consistent with each other [30]. The second one is the functional hierarchy index which is a measure of neuronal response to well-defined artificial stimuli. The Allen data provides several functional indexes [55], out of which we use the size of the receptive field (RF) that was named as the RF area measured with Gabor stimuli [55]. We also used the RF diameter that was measured with drifting sinusoidal gratings and provided in a separate study [30]. We calculated the means of the RF area from all the cells in the Allen data that show responses to Gabor stimuli. Since both studies have different numbers of cells, we used the mean values of both measures for six areas of the visual cortex. Totally we have four hierarchy indexes whose values are listed in Table S2.

## Supplemental Materials

**TABLE S1:** The number of cells and their brain areas in response to the movie stimulus.

| Brain Area | VISp | VISl | VISrl | VISal | VISpm | VISam | LGN | LP | APN | CA1 | DG | SUB | CA3 |
| --- | --- | --- | --- | --- | --- | --- | --- | --- | --- | --- | --- | --- | --- |
| Cell Units | 2015 | 933 | 1415 | 1553 | 879 | 1506 | 899 | 1448 | 667 | 2443 | 696 | 463 | 355 |

**TABLE S2:** The hierarchy across six areas of the visual cortex. Two anatomical hierarchy indexes (anatomical hierarchy score and hierarchical level) and two measures of receptive field (RF area and RF diameter) were taken from previous studies.

| Brain Area | V1 (VISp) | LM (VISl) | RL (VISrl) | AL (VISal) | PM (VISpm) | AM (VISam) |
| --- | --- | --- | --- | --- | --- | --- |
| Anatomical Hierarchy score [56] | -0.4210 | -0.0856 | -0.0549 | 0.0226 | 0.1124 | 0.2747 |
| Hierarchical level [30] | 0 | 0.4492 | 0.5035 | 0.6886 | 0.7031 | 0.8854 |
| RF area [55] | 589.23 | 716.83 | 646.50 | 713.86 | 787.43 | 769.63 |
| RF diameter [30] | 190.42 | 199.78 | 350.11 | 236.31 | 324.99 | 272.87 |

**Fig. S1:**
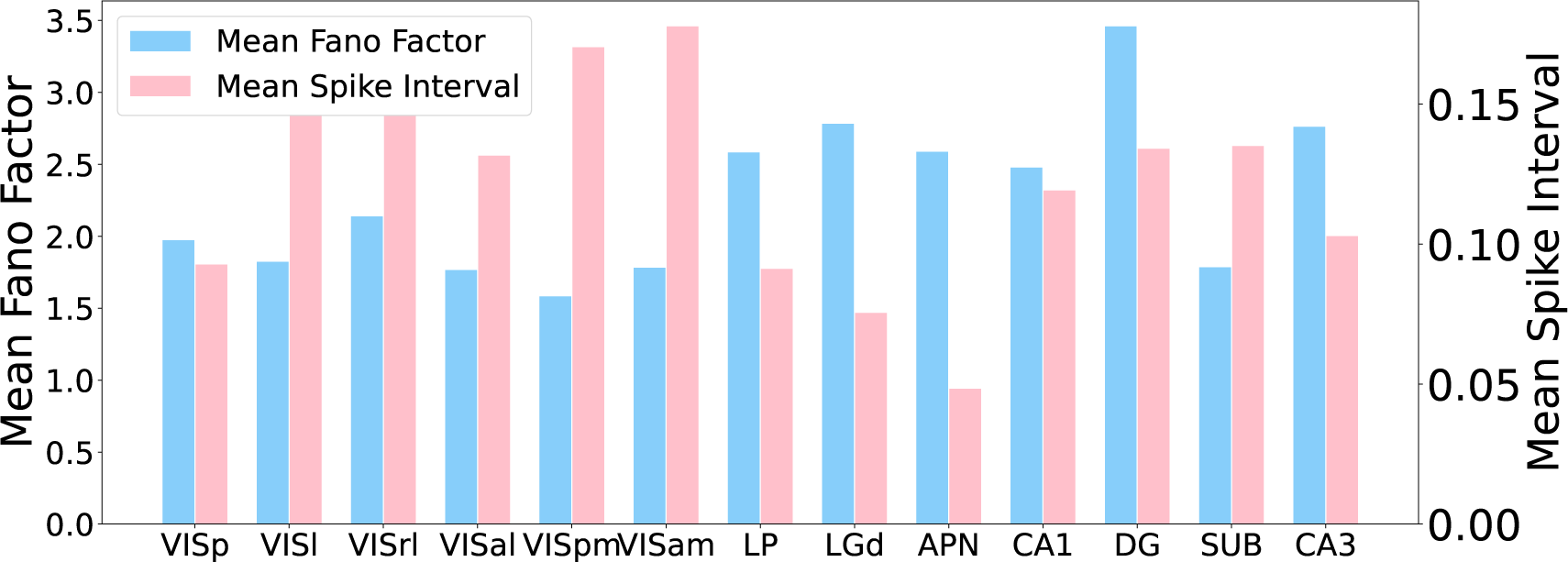
Related to Fig. 1E. The statics of firing activity in each brain. The distribution of inter-spike intervals and Fano factors.

**Fig. S2:**
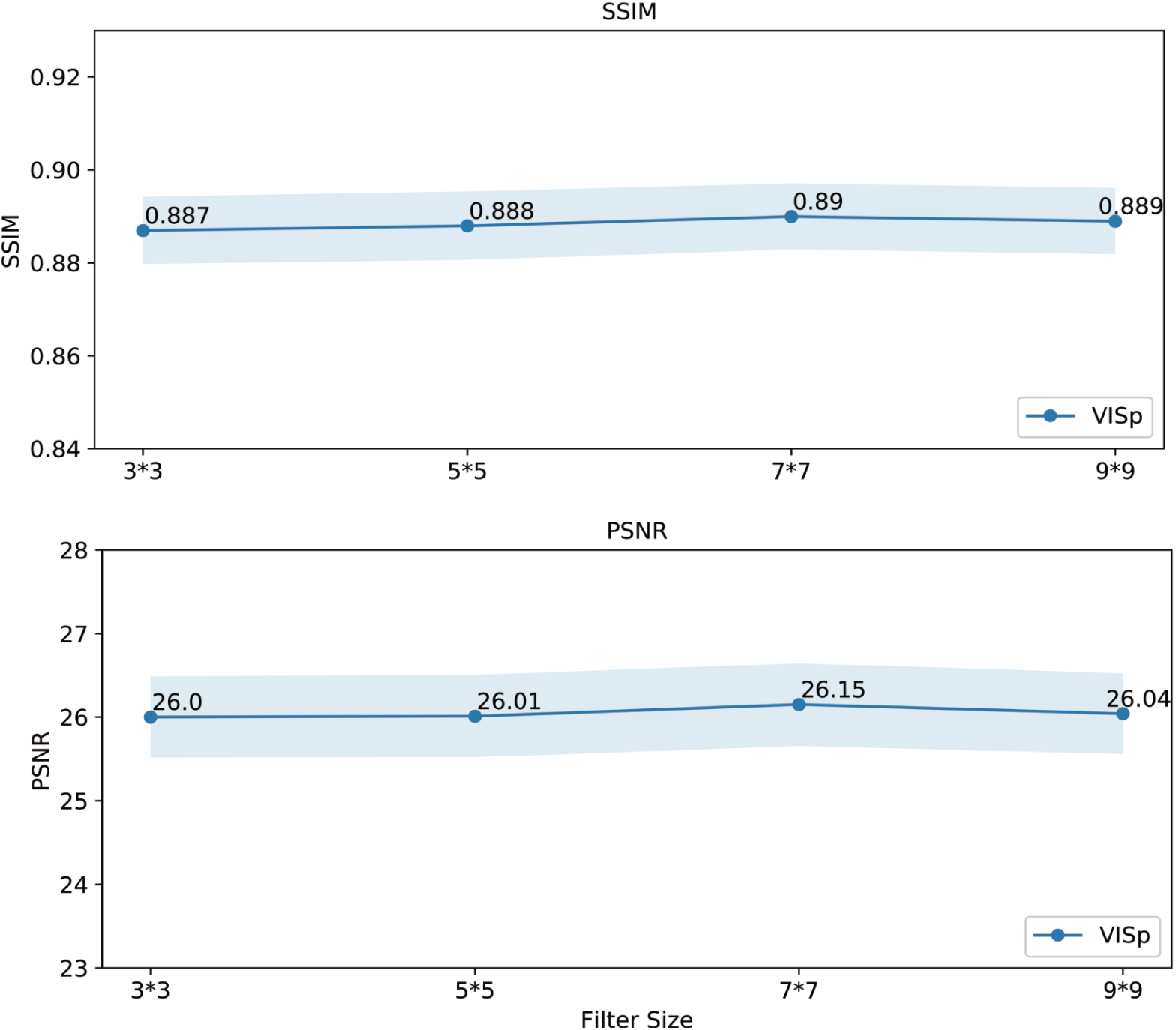
Related to Fig. 2. Decoding performance is robust to the variation of model parameter settings. Decoding metrics of VISp (using all 2015 cells) is robust to the change of the filter size in the model. The size of filters in the last few decoding layers of the network model was changed. The size used in the default model setting is 7*7.

**Fig. S3:**
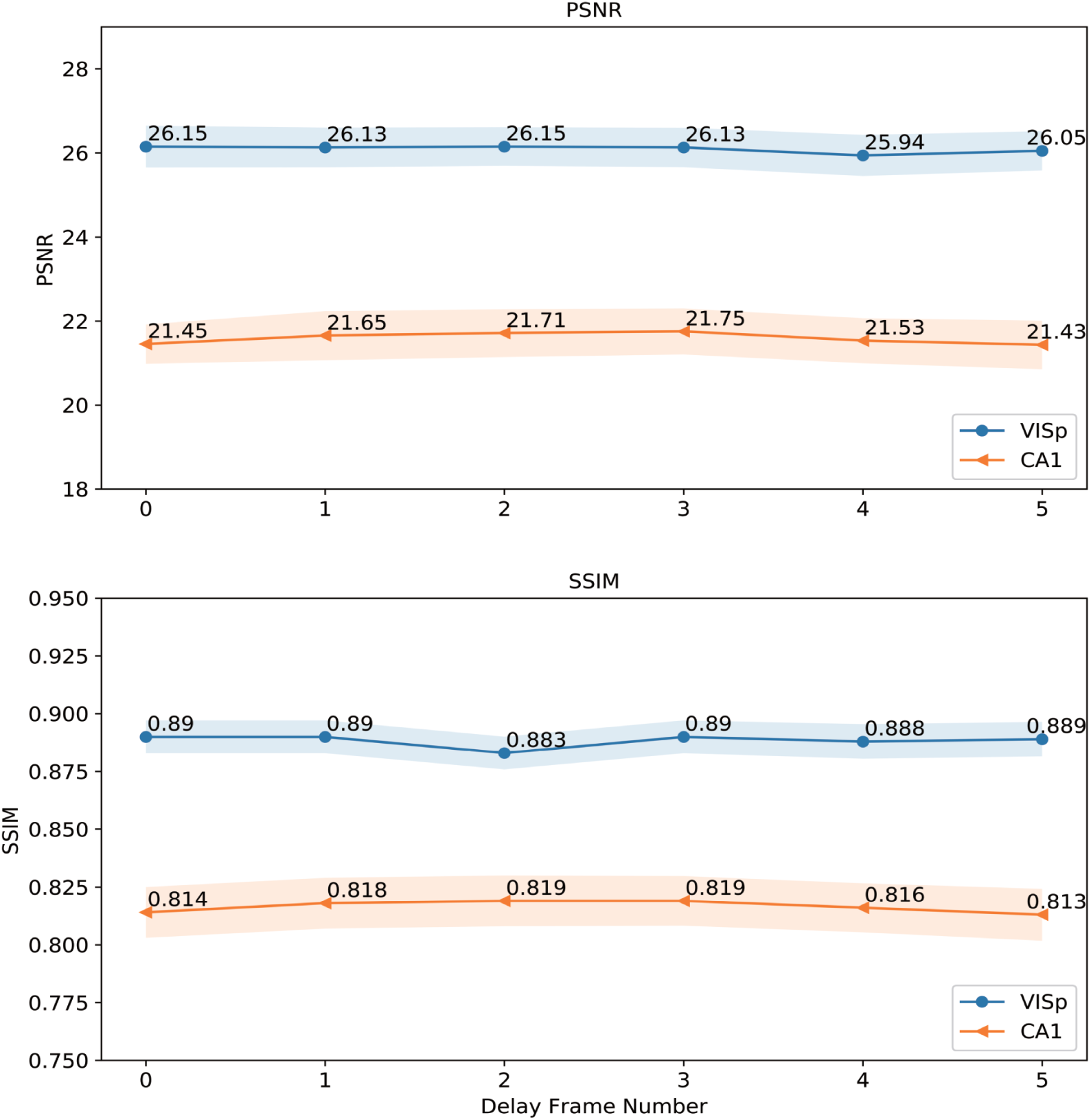
Decoding with time delays. Decoding results of VISp (using all 2015 cells) And CA1 (using all 2443 cells) with different time delays. Decoding was conducted by using the same neuronal response but with shifted video frame images different frames before the response time.

